# Ozone exposure upregulates the expression of host susceptibility protein TMPRSS2 to SARS-CoV-2

**DOI:** 10.1101/2020.11.10.377408

**Authors:** Thao Vo, Kshitiz Paudel, Ishita Choudhary, Sonika Patial, Yogesh Saini

## Abstract

**Background:** SARS-CoV-2, a novel coronavirus, and the etiologic agent for the current global health emergency, causes acute infection of the respiratory tract leading to severe disease and significant mortality. Ever since the start of SARS-CoV-2, also known as COVID-19 pandemic, countless uncertainties have been revolving around the pathogenesis and epidemiology of the SARS-CoV-2 infection. While air pollution has been shown to be strongly correlated to increased SARS-CoV-2 morbidity and mortality, whether environmental pollutants such as ground level ozone affects the susceptibility of individuals to SARS-CoV-2 is not yet established.

**Objective:** To investigate the impact of ozone inhalation on the expression levels of signatures associated with host susceptibility to SARS-CoV-2.

**Methods:** We analyzed lung tissues collected from mice that were sub-chronically exposed to air or 0.8ppm ozone for three weeks (4h/night, 5 nights/week), and analyzed the expression of signatures associated with host susceptibility to SARS-CoV-2.

**Results:** SARS-CoV-2 entry into the host cells requires proteolytic priming by the host-derived protease, transmembrane protease serine 2 (TMPRSS2). The TMPRSS2 protein and *Tmprss2* transcripts were significantly elevated in the extrapulmonary airways, parenchyma, and alveolar macrophages from ozone-exposed mice. A significant proportion of additional known SARS-CoV-2 host susceptibility genes were upregulated in alveolar macrophages and parenchyma from ozone-exposed mice.

**Conclusions:** Our data indicate that the unhealthy levels of ozone in the environment may predispose individuals to severe SARS-CoV-2 infection. Given the severity of this pandemic, and the challenges associated with direct testing of host-environment interactions in clinical settings, we believe that this mice-ozone-exposure based study informs the scientific community of the potentially detrimental effects of the ambient ozone levels determining the host susceptibility to SARS-CoV-2.

## Introduction

SARS-CoV-2, a novel coronavirus, and an etiologic agent of the current global health emergency, causes acute infection of the respiratory tract leading to severe disease and significant mortality ^1^. SARS-CoV-2 entry into the host cells is dependent upon the binding of the viral spike (S) protein to the host cellular receptor, angiotensin converting enzyme (ACE2), and its proteolytic priming by the host-derived protease, transmembrane protease serine 2 (TMPRSS2) ^2^. Therefore, the host susceptibility to SARS-CoV-2 could vary based on the expression of the host susceptibility proteins including ACE2 and TMPRSS2 ^3–5^. For example, increased expression of the host-derived protease, TMPRSS2, may promote the priming of SARS-CoV-2, thus resulting in increased infectivity and disease severity. The current literature indicates that individuals have varied susceptibility to SARS-CoV-2 that may be dependent on age ^6, 7^, gender ^8^, underlying comorbidities ^9^, and the environmental pollution ^10, 11^. However, the list of factors determining varied susceptibilities of human population to SARS-CoV-2 remains incomplete.

Nearly one-third of the United States population lives in areas with unhealthy levels of ozone ^12, 13^. While it is already known that the unhealthy levels of ozone increase the risk for developing cardiopulmonary health problems ^14–20^, it is unclear whether the ambient ozone levels regulate the expression of host susceptibility proteins to SARS-CoV-2 and, in turn, accounts for the varied susceptibilities of the human population to SARS-CoV-2. Addressing this critical question is highly relevant in terms of increasing our mechanistic understanding of the host-air pollution (environment) interactions underlying the SARS-CoV-2 pathogenesis and epidemiology, and for developing future preventative and therapeutic strategies.

To begin to understand the impact of ozone inhalation on the host susceptibility to SARS-CoV-2, we analyzed lung tissues collected from mice that were sub-chronically exposed to filtered-air or 0.8ppm ozone for three weeks (4h/night, 5 nights/week) ^21^, and analyzed the expression of gene and protein signatures associated with host susceptibility to SARS-CoV-2. We used western blotting and immunohistochemistry for assessing expression levels of TMPRSS2 protein in three different lung tissue compartments. To determine the RNA levels of Tmprss2 in a cell-specific manner, in situ gene expression was assessed for *Tmprss2* gene using RNAscope approach. Finally, the mRNA expression levels of 32 known SARS-CoV-2 host susceptibility genes were assessed in three lung tissue compartments using RNASeq approach. Our findings indicate that host-environment interaction may modulate the expression of host susceptibility proteins to SARS-CoV-2 and prime the host to manifest severe respiratory illness following SARS-CoV-2 infection.

## Methods

### Animal husbandry, experimental design and ozone exposure

Seven-week-old C57BL/6 mice were procured from Jackson Laboratory (Bar Harbor, ME). Mice were maintained in individually ventilated, hot-washed cages on a 12-hour dark/light cycle. Mice were housed in polycarbonate cages and fed a regular diet and water *ad libitum*. All animal use procedures were performed after approval from the Institutional Animal Care and Use Committee (IACUC) of the Louisiana State University. Ozone was generated by ozone generator (TSE Systems, Chesterfield, MO), and the ozone levels were monitored by UV photometric ozone analyzer (Envia Altech Environment, Geneva, IL). Data acquisition was done through DACO monitoring software (TSE Systems, Chesterfield, MO). Control mice were kept in chamber supplied by filtered air (Air). Animals were exposed to ozone (800ppb; 4h/night, 5 nights/week, for 3 weeks) or air. Further details have been published previously ^1^.

### Necropsy and tissue harvesting, micro-dissection of the extrapulmonary airways, the parenchyma, and Purification of airspace macrophages

Animals were euthanized and tissues were collected for RNA isolation or histological analyses, as described previously ^1^.

### Immunohistochemistry for TMPRSS2

Formalin-fixed, paraffin-embedded 5μm lung sections were used for immunohistochemical localization of TMPRSS2. Sections were deparaffinized with Xylene and rehydrated with graded ethanol. Heat-induced antigen-retrieval was performed using a Citrate buffer (pH 6.0). Endogenous peroxidases were quenched with 3% hydrogen peroxide (10 min at room temperature). After blocking with 3% goat serum for 30 min, sections were incubated for 2h at room temperature with rabbit polyclonal TMPRSS2 primary antibody (ab214462; Abcam Cambridge, MA). The sections were then processed using VECTASTAIN Elite ABC HRP Kit (Vector Laboratories, Burlingame, CA), followed by chromogenic substrate conversion to insoluble colored precipitate using ImmPACT NovaRED HRP substrate Kit (Vector Laboratories, Burlingame, CA). Sections were counterstained with Gill’s Hematoxylin-I, dehydrated, and coverslipped with mounting media (H-5000, Vector Laboratories, Burlingame, CA).

### Western Blotting for TMPRSS2

Whole lung homogenates and bronchoalveolar lavage aliquots were separated by SDS-PAGE (NuPAGE 4-12% Bis-Tris gradient gel; Life Technologies, CA) and transferred to PVDF membrane. Rabbit polyclonal TMPRSS2 primary antibody (ab214462; Abcam Cambridge, MA) and mouse monoclonal alpha tubulin (T5168, Sigma-Aldrich, MO) were used. Protein bands were visualized using secondary antibodies (alexa fluor 680 Goat anti-rabbit IgG or Alexa fluor 800 Goat anti-mouse IgG) and acquired using Odyssey CLx, Imager (LI-COR, NE) ^2^.

### *In situ* localization of *Tmprss2* mRNA

Formalin-fixed paraffin-embedded 5μm lung sections were used for *in situ* localization of *Tmprss2* mRNA using RNAscope technologies, as reported previously ^1, 3^.

### RNA isolation and quality assessment, Construction of sequencing library, RNA sequencing and Gene Expression Analyses, and Data Availability

The detailed methodologies have been previously published ^1^. The raw data have been submitted to the Gene Expression Omnibus (GEO) database. The data is available via https://www.ncbi.nlm.nih.gov/geo/query/acc.cgi?acc=GSE156799.

### Statistical analyses

Student’s T test for was used to determine significant differences among groups. Statistical analyses were performed using GraphPad Prism 8.0 (GraphPad Software, La Jolla, CA). All data were expressed as mean ± standard error of the mean (SEM). A *p*-value<0.05 was considered statistically significant.

## Results and discussion

TMPRSS2 is essential for the proteolytic priming of viral spike (S) protein of the SARS-CoV-2 following its binding to host receptor, ACE2. In fact, a recent study elegantly demonstrated that host cell entry of SARS-CoV-2 can be blocked by a clinically proven inhibitor of TMPRSS2 indicating the critical importance of TMPRSS2 in determining SARS-CoV-2 infectivity ^2^. Therefore, the host susceptibility to SARS-CoV-2 could vary based on the expression of the host susceptibility proteins including ACE2 and TMPRSS2 ^3–5^. Individuals have varied susceptibility to SARS-CoV-2 that may be dependent on various factors including air pollution ^10, 11^. Air-pollution levels correlate strongly with increased morbidity and mortality due to SARS-CoV-2 ^22–24^. While it is already known that the unhealthy levels of ozone, one of the 6 criteria air pollutants, increase the risk for developing cardiopulmonary health problems ^14–20^, it is unclear whether the ambient ozone regulates the levels of expression of host susceptibility proteins to SARS-CoV-2 and in turn accounts, in part, for the varied susceptibilities of human population to SARS-CoV-2.Therefore, we sought to test the hypothesis that ozone induces the expression of TMPRSS2 in lung tissue.

As compared to filtered air-exposed mice, the ozone-exposed mice had significantly elevated levels of TMPRSS2 protein in the whole lung lysate (**Fig 1A**) and the bronchoalveolar lavage fluid (BALF) (**Fig 1B**). These results demonstrate that ozone exposure increases the expression of TMPRSS2 protein in the lungs of mice and that this protein is detectable in the BALF fluid. Next, we performed immunohistochemical staining on lung sections to visualize cell-specific localization of TMPRSS2. Interestingly, while the TMPRSS2 staining was evident in the airway epithelium and alveolar macrophages from filtered air-exposed mice, the staining intensity was remarkably increased in the airway epithelial cells, alveolar epithelial cells, and alveolar macrophages from ozone-exposed mice (**Fig 1C**). These results clearly demonstrate that ozone increases the expression of TMPRSS2 in the lung tissue in a cell-specific manner.

**Figure 1.**
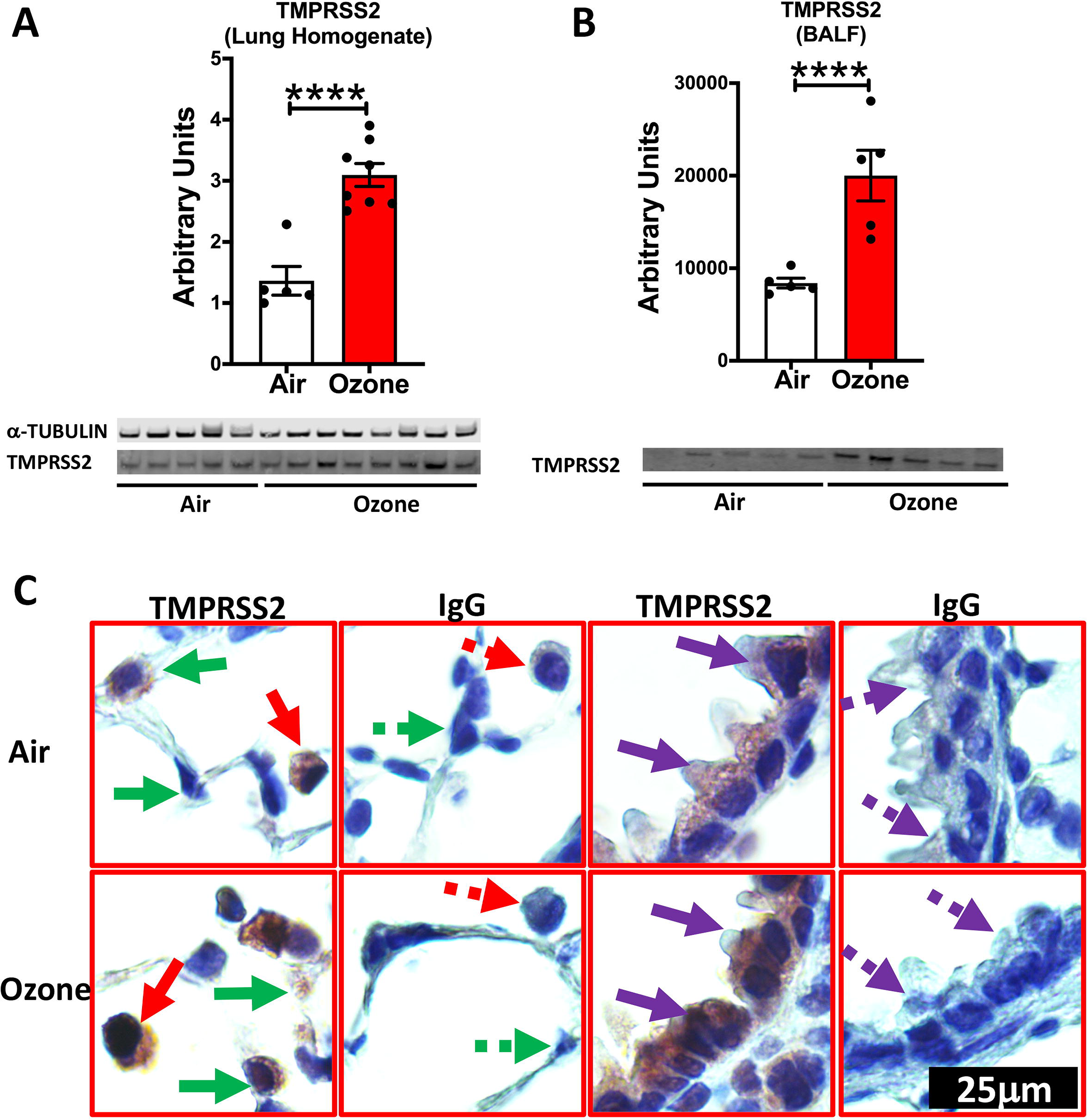
TMPRSS2 protein expression is upregulated in the lungs of ozone exposed mice: **(A)** Western blot analyses [Bar Graph (above), and representative gel image (below) showing bands for TMPRSS2 protein and Tubulin loading control] on the whole lung homogenate from Air- and Ozone-exposed mice (n=6-8). **(B)** Western blot analyses [Bar Graph (above), and representative gel image (below) showing bands for TMPRSS2 protein] on the equal volumes (loading control) of bronchoalveolar lavage fluid from Air- and Ozone-exposed mice (n=5). The data are expressed as means (±SEM). Student’s T-Test **** *P*<0.0001. **(C)** Immunohistochemical staining for TMPRSS2 in macrophages (solid red arrow), alveolar epithelial cells (solid green arrow), and bronchiolar epithelial cells (solid Purple arrow). Negatively stained cells are indicated by dotted arrows in lung sections that were incubated with antibody (IgG) control. All images were captured at the same magnification.

In order to test whether TMPRSS2 protein expression correlates with mRNA expression in a cell-specific manner, we analyzed RNASeq dataset from extrapulmonary airways (**Fig. 2A**), parenchyma (**Fig. 2B**), and alveolar macrophages from filtered-air and ozone exposed mice (**Fig. 2C**) for *Tmprss2* transcript levels. As expected, the fragment per kilo base per million mapped reads (FPKM) for *Tmprss2* were significantly upregulated in all the three tissue compartments in ozone-exposed as compared to filtered-air exposed mice. We further confirmed these findings for cell-specificity in ozone-exposed airways and alveoli using RNAscope-based in situ hybridization. This assay also showed significantly increased signals for *Tmprss2* transcripts in both the airway epithelial cells and the alveolar epithelial cells of ozone-exposed compared to filtered-air exposed mice (**Fig. 2D**). These data suggest that the changes in the protein levels of TMPRSS2 are a result of changes at the level of gene expression indicating that ozone directly or indirectly differentially regulates the gene expression of *Tmprss2*.

**Figure 2.**
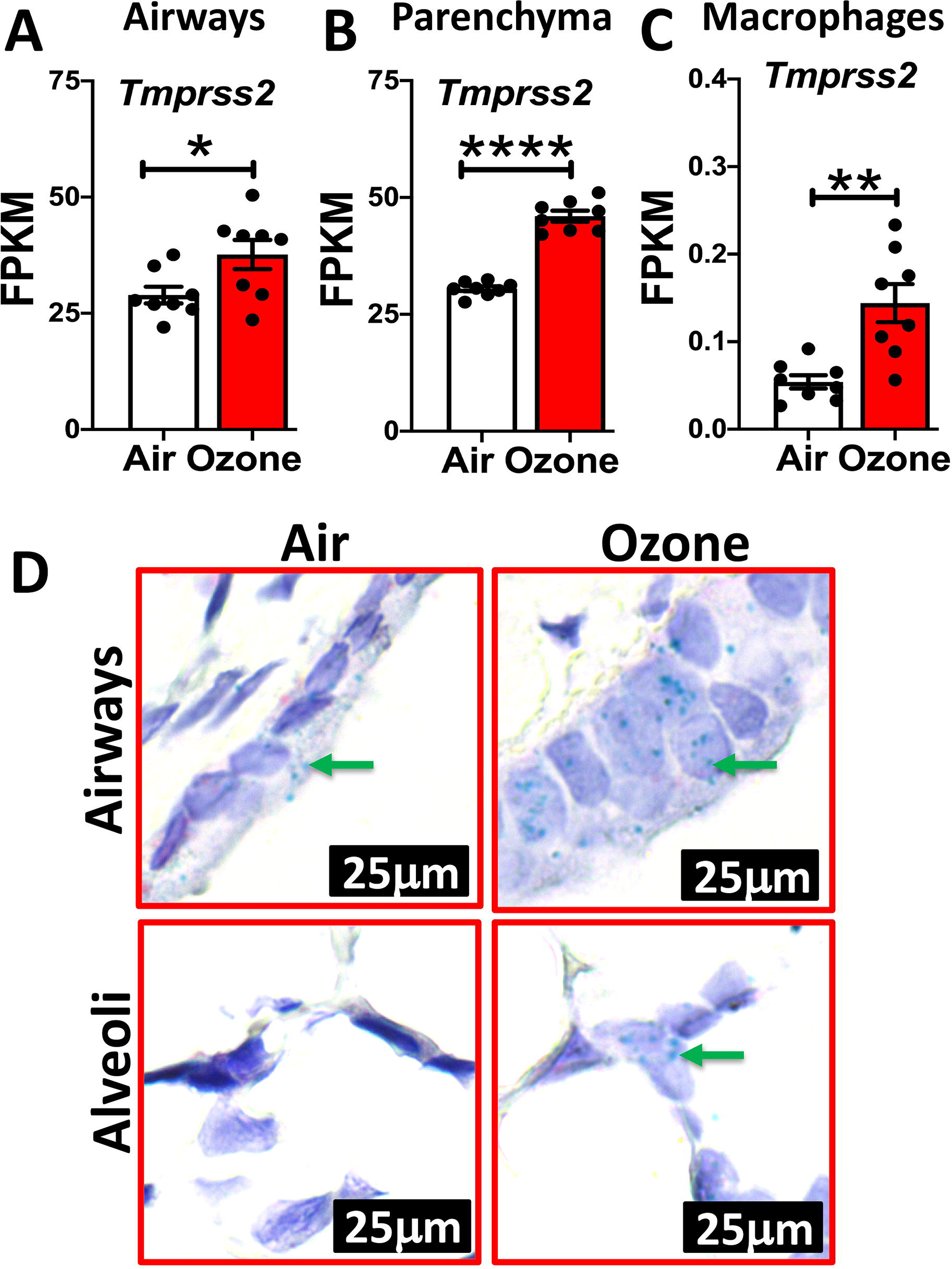
*Tmprss2* mRNA expression is upregulated in the lungs of ozone exposed mice: FPKM values obtained from RNAseq data set obtained from airways **(A)**, parenchyma **(B)**, and alveolar macrophages **(C)** were used to quantify relative expression levels of Tmprss2 mRNA in air-versus ozone-exposed mice (n=8). The data are expressed as means (±SEM). Student’s T-Test **** *P*<0.0001. **(D)** RNAscope-based in situ hybridization for *Tmprss2* transcripts (green dots representing punctate staining for *Tmprss2* mRNA in airway epithelial cells (Top) and alveolar epithelial cells (bottom) in air-(left) and ozone-exposed (right) mice. All images were captured at the same magnification.

Finally, from the RNASeq data, we compared the normalized z-scores of 32 known genes associated with host susceptibility to SARS-CoV-2. Of the three tissues analyzed, while only four of the total 32 genes, i.e., *Furin, Thop1, Ppia, and Tmprss2*, were significantly increased in the extrapulmonary airways of ozone-exposed versus filtered-air exposed mice (**Fig. 3A**); in the lung parenchyma, nearly half of the host susceptibility genes were upregulated while the rest half were downregulated in ozone-exposed versus filtered-air exposed mice (**Fig. 3B**). In contrast to the extrapulmonary airways and the lung parenchyma, a major proportion of the host susceptibility genes including those of *Tmprss2, Ace, Anpep, Cd4*, and *Ccr5* were significantly upregulated in the CD11b^−^ lung macrophages from ozone-exposed versus filtered-air exposed mice (**Fig. 3C**) indicating a large effect of ozone-exposure on SARS-CoV-2 host susceptibility genes in CD11b^−^ lung macrophages

**Figure 3.**
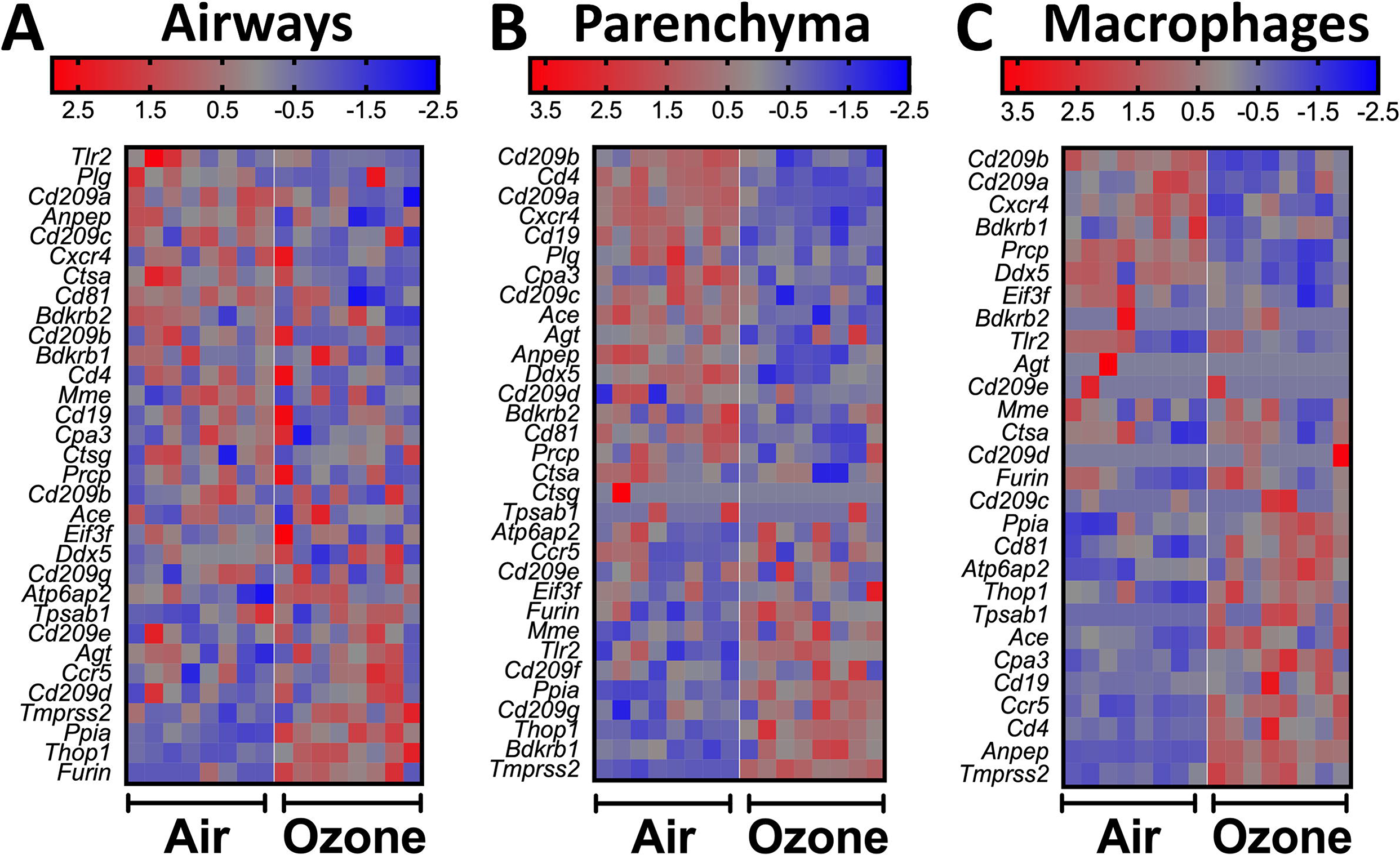
Heatmap depicting normalized gene expression values (Z-scores) of genes associated with SARS-CoV-2 host susceptibility in airways **(A)**, parenchyma **(B)**, and alveolar macrophages **(C)**from air- and ozone-exposed mice.

In conclusion, this study suggests a critical role of the host-environment (ozone pollution) interaction in modulating the susceptibility of the human population to SARS-CoV-2. Since TMPRSS2 is essential for the proteolytic processing of several coronaviruses including SARS-CoV-1, MERS, as well as influenza A virus ^25, 26^, our findings have implications beyond SARS-CoV-2 infections. Taken together, this study presents a novel finding that will have significant and immediate impact on our understanding of the pathogenesis and epidemiology of SARS-CoV-2 pandemic.

## Acknowledgments

We thank Sherry Ring for assistance with histological tissue processing.

## Author Contributions

Y.S. and S.P. conceived and designed the research; T.V. performed the RNA in situ hybridization assays; K.P. performed the immunohistochemical staining; I.C. performed immunoblotting assays. Y. S., T.V., K.P., and I.C., maintained the animal colony, performed ozone-exposures, and conducted animal necropsies. S.P. and Y.S. analyzed the histopathological, immunohistochemical, and RNA in situ hybridization data and wrote and reviewed the manuscript for intellectual contents.

